# A data harmonization pipeline to leverage external controls and boost power in GWAS

**DOI:** 10.1101/2020.11.30.405415

**Authors:** Danfeng Chen, Katherine Tashman, Duncan S. Palmer, Benjamin Neale, Kathryn Roeder, Alex Bloemendal, Claire Churchhouse, Zheng Tracy Ke

## Abstract

The use of external controls in genome-wide association study (GWAS) can significantly increase the size and diversity of the control sample, enabling high-resolution ancestry matching and enhancing the power to detect association signals. However, the aggregation of controls from multiple sources is challenging due to batch effects, difficulty in identifying genotyping errors, and the use of different genotyping platforms. These obstacles have impeded the use of external controls in GWAS and can lead to spurious results if not carefully addressed. We propose a unified data harmonization pipeline that includes an iterative approach to quality control (QC) and imputation, implemented before and after merging cohorts and arrays. We apply this harmonization pipeline to aggregate 27,517 European control samples from 16 collections within dbGaP. We leverage these harmonized controls to conduct a GWAS of Crohn’s disease. We demonstrate a boost in power over using the cohort samples alone, and that our procedure results in summary statistics free of any significant batch effects. This harmonization pipeline for aggregating genotype data from multiple sources can also serve other applications where individual level genotypes, rather than summary statistics, are required.

## 1. Introduction

Genome-wide association studies (GWAS) have been successful in identifying genetic loci that confer some risk to disease. ^1,2,3,4,5,6^ A key factor that determines the ability of GWAS to detect disease-associated variants is sample size. Leveraging external controls that have already been genotyped and shared publicly can increase power for discovery while allowing resources to be focused on collecting and genotyping only cases.^7^ Integrating external control samples can also supplement existing controls and increase the number of ancestrally matched controls in a study.

Despite these clear advantages, external control resources have not been widely adopted. One reason is the significant administrative and technical barriers to obtaining permission and then acquiring multiple different publicly available data sets.^8,9^ Furthermore, it is critical to ensure that any allele frequency differences between controls and cases are indeed attributable to the disease or trait being studied, and not due to systematic biases caused by insufficient ancestry matching, technical artifacts, or batch effects.

The use of external controls typically requires merging genotype data from multiple sources in order to maximize control sample size and provide a large pool of controls from which to match the ancestry of the cases. In this process, many factors such as genotyping error, batch effects, and the use of different genotyping platforms can yield spurious correlations between cases and controls^10^. As such, it is crucial to conduct careful quality control and data harmonization that is targeted towards these different sources of error when merging external controls. Even when genotyping data is derived from a single cohort, its use in GWAS first requires some analyses to ensure the data is of sufficient quality to be employed in association testing. This quality control (QC) usually involves a series of analytic filters aimed at removing poor quality samples and problematic single nucleotide polymorphisms (SNPs).

To this aim, we have developed a data harmonization pipeline to reliably leverage external genotyped controls, aggregated from multiple sources, to boost power in GWAS without introducing spurious associations. The design of the pipeline addresses two key issues in merging such heterogeneous data - one is errors driven by batch effects, and the other is spurious signals introduced by imputation. Our pipeline iterates through a series of QC filters and imputation to examine samples at the levels of cohort and genotyping platform. We demonstrate the utility of our harmonization pipeline by aggregating 27,517 European control samples from 16 data collections within dbGaP,^11,12^ and use these as controls in a GWAS of Crohn’s disease. Our work demonstrates the plausibility of harmonizing genotyping data from multiple sources, and enables external controls to be reliably integrated in GWAS.

The application of our data harmonization pipeline is not limited to GWAS, but may be useful for many other analysis methods that require individual-level genotype data rather than summary statistics. One example is the knockoff method for controlling false discovery rates^13,14^ and its application to genetic association studies. This method creates surrogate genes (“knockoff variables”) from individual-level genotypes, handling linkage disequilibrium in a principled manner.^15^ Another type of analysis utilizes the genetic relationship matrix to capture the pairwise relationship among individuals to compute estimates of heritability, the genetic correlation between traits, and genetic risk scores.^16,17^ When computing genetic risk scores for highly polygenic traits such as autism spectrum disorder, for which GWAS discoveries have been limited, this approach is more sensitive than traditional polygenic risk score approaches that instead utilize summary statistics.^18^ Despite the tremendous progress leveraging summary statistics, many valuable avenues of analysis require the aggregation of genotypes, thus a reliable harmonization pipeline is essential to remove batch effects. Although we demonstrate the performance of our pipeline in the context of GWAS, the proposed data harmonization method is also promising for these other applications.

## 2. Material and Methods

### 2.1. The data harmonization pipeline

The pipeline contains four modules: (i) Within-array processing, (ii) Imputation, (iii) Cross-array comparison, and (iv) Re-imputation. The *Within-array processing* module aims to group samples by array, cohort, and ancestry, so that each group contains homogeneous samples. Within this module, multiple QC filters are applied to samples and variants, within and across cohort. Next, *Imputation* is conducted in each homogeneous group. The post-imputation data are merged within each array, followed by a few standard QC filters. The first two modules resolve issues such as genotyping errors and missing values; however, two issues remain. First, batch effects still exist, which prohibit us from merging data across different arrays. The *Cross-array comparison* module detects batch effects via “pseudo-GWAS”, where samples from one array are treated as cases and samples from the other arrays are treated as controls. Significant variants in this pseudo case-control comparison will be removed. Second, the imputation quality is low for some variants, possibly due to low coverage in the reference panel or high recombination rate. Such low-quality imputation can drive false association signals in GWAS. In the *Re-imputation* module, we aim to detect such spurious signals and remove them, before re-imputing in the surrounding region to recover those QC-failed variants and improve imputation quality. A summary of the data harmonization pipeline is shown in Figure 1. Below, we give a high-level description of each module. The details can be found in Appendix B.

**Figure 1.**
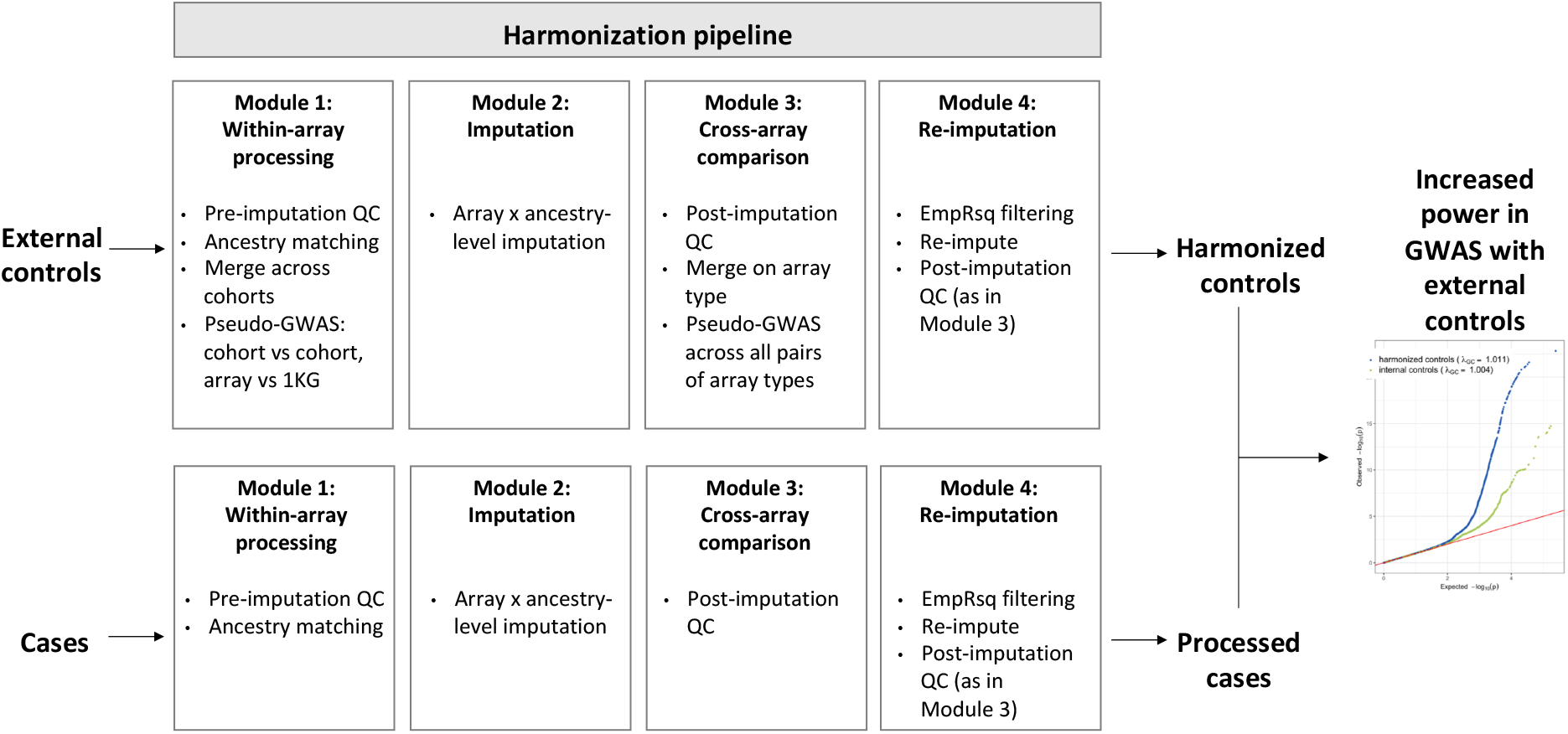
A high-level description of the data harmonization pipeline.

#### Module 1: Within-array processing

consists of the following main steps:

- *Cohort-level QC*. Remove samples with high missingness rate and abnormal in-breeding coefficients, as well as variants with a high missingness rate.
- *Ancestry matching*. Each sample is assigned to one of the five ancestries, East Asian, South Asian, African, American, and European. Here, we focus on European ancestry, where samples are further split into three sub-groups - major European, Finnish, and Ashkenazi Jewish. The ancestry assignment is determined by a pre-trained classifier, where the training data is 1000 Genomes data (with self-reported ancestry labels) and the classification method is a standard random forest algorithm^19^ on leading PC’s.
- *Merging*. Based on the first two steps, a stratum has been formed for each array-cohort-ancestry combination. For each stratum, we further remove variants with low minor allele frequencies and small Hardy-Weinberg Equilibrium p-values. We then merge samples of the same genotyping array and ancestry group.
- *Array-level QC*. To remove batch effects, two rounds of pseudo-GWAS are performed iteratively within each array-ancestry stratum. In the first round, all the samples in this stratum are compared with the 1000 Genomes samples belonging to the same ancestry group. In the second round, the samples from one cohort are compared with those from the other cohorts in the same array-ancestry stratum. In these pseudo-GWAS comparisons, the first 20 principal components (PCs) are included as covariates to account for population structure. Significant variants identified in either round are removed.

#### Module 2: Imputation

Module 1 produces a data stratum per array per ancestry. These strata are imputed separately. The motivation is that each stratum contains relatively homogeneous samples, which can improve the imputation quality compared with imputing all strata together. We use the Michigan Imputation server with 1000 Genomes data as the reference panel.

#### Module 3: Cross-array comparison

While Module 1 (through the *array-level QC*) is aimed at accounting for batch effects due to genotyping across independent studies, we include a second module to target batch effects that arise due to imputation. We expect that the quality of imputation at different regions of the genome will vary for different arrays, owing to the design of their particular backbone and to technical characteristics. This will induce considerable batch effects due to the large number of variants that are imputed in Module 2 (about 46.8M per stratum in our experiment). This module aims to remove such batch effects by cross-array pseudo-GWAS.

- *Post-imputation QC*. Remove variants with low minor allele frequencies, or small Hardy-Weinberg Equilibrium p-values, or low imputation info scores.
- *Cross-array pseudo-GWAS*. We first merge samples genotyped on the same array type through an “inner join” operation. Next, a pseudo-GWAS is performed for each array, where samples on this array are treated as cases and those on all other arrays are used as controls. Since ancestry groups have been merged, we include 20 leading PC’s as covariates in the pseudo case-control comparison, to account for cross-ancestry heterogeneity. Significant variants are removed.

#### Module 4: Re-imputation

The last module deals with spurious association signals introduced by poor quality imputation, first removing the poorly typed variants that drive the imputation and then re-imputing the surrounding region.

- *Filtering*. There are multiple metrics to assess imputation quality. ^20,21,22,23^ We use EmpRsq, which is the squared correlation between the leave-one-out imputed dosages and the observed (i.e. typed) genotypes. Any typed variant with EmpRsq below the minimum threshold is removed.
- *Re-imputation*. With those poorly typed variants already removed, we re-do the whole imputation procedure, where we use the same strata as in Module 1, conduct imputation in a similar way as in Module 2, and then apply similar post-imputation steps as in Module 3.

The strength of this pipeline comes from the division of QC procedures into steps that are aimed at capturing genotyping batch effects and those that are designed to identify bad sites arising from poor imputation. The order of modules in the pipeline makes this possible, and the cross-array pseudo-GWAS ensures that the allele frequencies are consistent between chips.

The *Re-imputation* module plays a key role in removing poorly genotyped SNPs that pass earlier QC filters but show evidence of driving low quality inference at nearby imputed sites. The first, rather strict, filter aims to ensure that the retained (typed) variants are of high quality. We then conduct a second imputation, so that the poorly imputed variants in the previous round are corrected. Subsequent QC of the re-imputed data indeed removes much less variants, confirming the benefit of the re-imputation module.

When this pipeline is used to harmonize external control data, the same process should be applied to the case samples. We have constructed our data harmonization pipeline as a series of modules containing multiple filters, the thresholds and parameters of which may be adjusted by the user. We have selected their values to be effective in our applications. The code to execute the pipeline is shared publicly in our GitHub repository (https://github.com/mikkoch/unicorn-qc).

## 3. Results

### 3.1. Creating a resource of harmonized external controls

We aggregated genotyped data on 27,517 individuals of self-reported European-descent from 16 studies in the dbGaP repository. These cohorts had been genotyped using a plethora of technologies including various Illumina and Affymetrix arrays. A summary of external controls is in Table 1, and a detailed description of 16 cohorts is provied in Table 2 of Appendix A.

**Table 1.** Summary of controls from 16 genotyping studies, grouped by array type. For each array, the number of SNPs after harmonization includes the imputed ones. The number of SNPs in the combined data set is obtained by an inner join.

| Genotyping Array | #Cohorts | Before harmonization |  | After harmonization |  |
| --- | --- | --- | --- | --- | --- |
|  |  | #Samples | #SNPs | #Samples | #SNPs |
| Affy6 | 3 | 4,504 | 907k | 2,936 | 6.18M |
| Axiom | 1 | 10,000 | 636k | 9,080 | 5.73M |
| Human300 | 1 | 219 | 300k | 197 | 6.42M |
| Human550 | 5 | 5,559 | 550k | 4,449 | 6.33M |
| Human610 | 4 | 4,672 | 610k | 3,604 | 6.19M |
| Human660 | 2 | 2,563 | 660k | 2,244 | 6.25M |
| Combined | 16 | 27,517 | - | 22,510 | 5.52M |

**Table 2.** The 16 studies used for creating a resource of external controls, taken from the dpGAP repository.

| Study | dbGaP Accession Number | Ancestry | #Samples | Genotyping Chip |
| --- | --- | --- | --- | --- |
| Late Onset Alzheimer’s Disease Family Study: Genome-Wide Association Study for Susceptibility Loci | phs000168.v2.p2 | European | 1292 | HumanHap 610 |
| GENEVA Genes and Environment Initiatives in Type 2 Diabetes | phs000091.v2.p1 | European | 3148 | Affymetrix 6.0 |
| National Eye Institute Glaucoma Human Genetics Collaboration (NEIGHBOR) Consortium Glaucoma Genome-Wide Association Study | phs000238.v1.p1 | European | 1240 | HumanHap 610 |
| A Study of the Genetic Causes of Complex Pediatric Disorders | phs000490.v1.p1 | European | 2631 | HumanHap 610/<br>HumanHap 550 |
| Genome-Wide Association Study of Schizophrenia | phs000021.v3.p2 | African/<br>European | 2332 | Affymetrix 6.0 |
| Neurodevelopmental Genomics: Trajectories of Complex Phenotypes | phs000607.v2.p2 | European | 5715 | HumanHap 610/<br>HumanHap 550 |
| Sweden-Schizophrenia Population-Based Case-Control Exome Sequencing | phs000473.v2.p2 | European | 773 | HumanHap550 |
| National Human Genome Research Institute (NHGRI) GENEVA Genome-Wide Association Study of Venous Thrombosis | phs000289.v2.p1 | European | 1310 | HumanHap 610 |
| Whole Genome Scan for Pancreatic Cancer Risk in the Pancreatic Cancer Cohort Consortium and Pancreatic Cancer Case-Control Consortium (PanScan) | phs000206.v5.p3 | European | 4133 | HumanHap 610/<br>HumanHap 550 |
| Research Program on Genes, Environment and Health (RPGEH) | phs000788.v1.p2 | European | 10000 | Axiom KP |
| NIH Exome Sequencing of Familial Amyotrophic Lateral Sclerosis Project | phs000101.v4.p1 | European | 773 | HumanHap 550 |

We applied the harmonization pipeline. Module 1 removed 5,007 samples, where 1,570 were filtered due to removal of non-European samples (the majority of non-European samples are from the GERA cohort, while other cohorts contain very few non-European individuals), 579 due to high sample missing rate, and 2,858 because of family relatedness or abnormal inbreeding coefficients. A total of 22,510 samples were retained. The QC steps in Modules 2-4 only removed variants but not samples. In additional to standard QC filters (missing rate, minor allele frequencies and Hardy-Weinberg Equilibrium) the cross-array pseudo-GWAS removes around 500 variants on average from each genotyping platform. We merged across the different platforms, retaining any SNPs with missing rate < 1%, to obtain the final harmonized dataset of 5,524,462 variants.

Figure 2 shows the projection of the harmonized controls onto the first two PC’s of the 1000 Genomes (1KG) data and onto the first two PCs of its European subset. The left panel shows that the majority of harmonized controls are indeed of European descent, and the right panel shows that they represent a range of European ancestry, including British, Italian, Northern European and Spanish, and are distinct from Finland. The set of harmonized public controls provide a much richer resource of European genotypes as it is approximately ten times larger than the European subset of 1KG and represents a more continuous sample along the cline of European ancestry. While this particular dataset was developed in the first instance as a European ancestry control resource, due to the availability of larger sample numbers, the same pipeline may be applied to studies of other ancestries to generate harmonized controls for more broad collection of genome-wide association studies. A list of SNPs that were removed in the harmonization pipeline, along with details of the step in which each was filtered out, is available in our GitHub repository (https://github.com/mikkoch/unicorn-qc).

**Figure 2.**
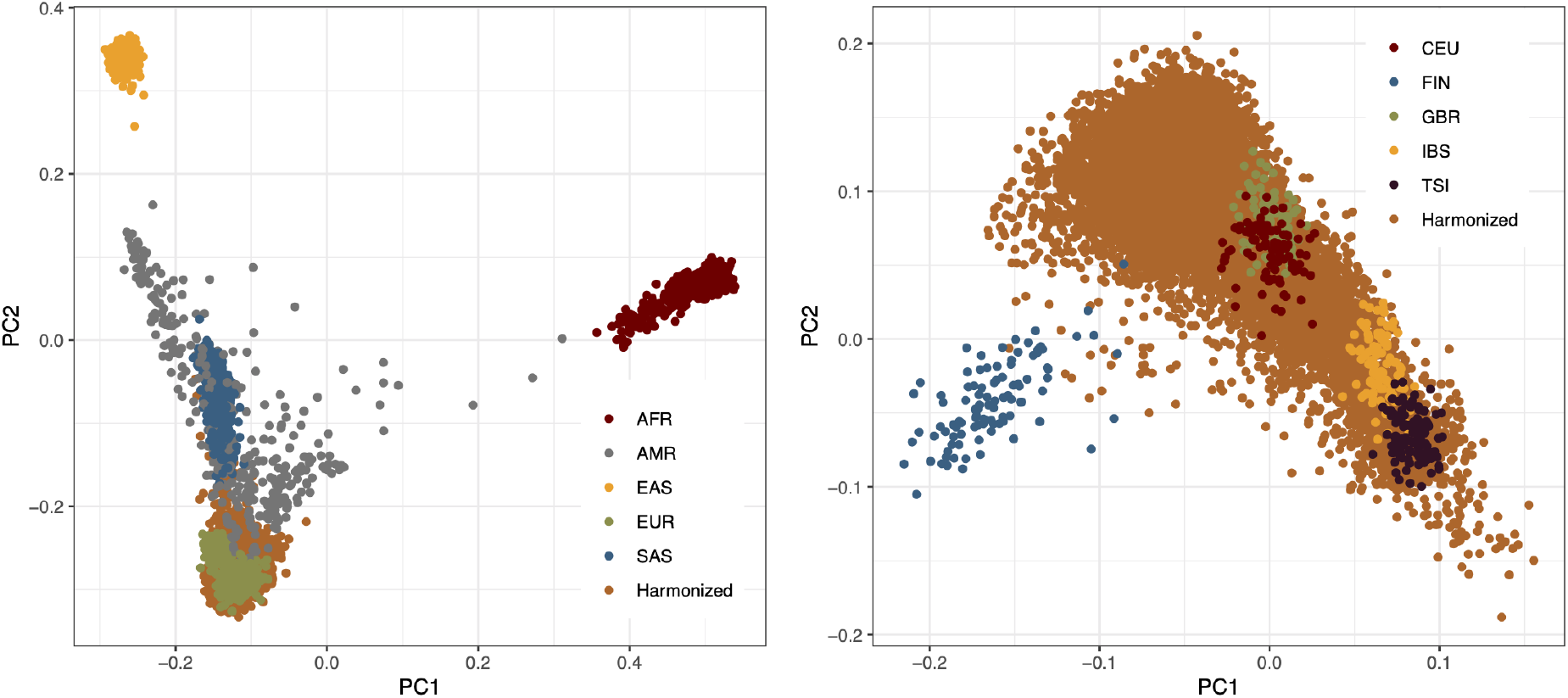
Principal component (PC) analysis representation of the ancestry distribution of the harmonized public controls. Left: projection of harmonized controls onto the PC space of 1000 Genomes (1KG). Right: projection onto the PC space of 1KG European subset.

### 3.2. Performance of GWAS using the external control resource

To access the quality and utility of the harmonized public control data set, we conducted a comparison of GWAS using the Crohn’s disease data of the CHOP study,^24^ obtained from the IBD Genetics Consortium. This study collected 1,589 cases and 5,950 internal controls. We examined p-values calculated from comparing the IBD cases to (i) the harmonized controls with those obtained from using (ii) the IBD study *internal* controls, as well as (iii) p-values from the IBD consortium’s meta-analysis, which included 5,956 case subjects and 14,927 control subjects^24^. The meta-analysis results serve as our best picture of the truth. We applied the same harmonization pipeline to the Crohn’s disease data, but Module 3 (the cross-array comparison) is not required as all of the IBD study data is typed on a single chip (see Materials and Methods).

We consider two performance characteristics: (a) a boost in power for true signals and (b) the elimination of spurious signals. We compare the association results generated from the harmonized controls and those obtained from using the internal controls in the Manhattan plots of Figure 3 and see that in general they are highly concordant across the whole genome. The QQ plot in the bottom right panel shows that there is very little evidence of inflation in either version of the GWAS (λ_*GC*_ = 1.011 using harmonized controls and 1.004 using internal controls). The QQ plot also shows the increase in significance of the most highly associated SNPs when using the harmonized controls compared to the internal controls. The bottom left panels show a zoomed-in view of two regions of genome-wide significant signal in the meta-analysis (on chromosomes 3 and 5) where, even at this finer scale, the concordance is still very good. Furthermore, the minimum p-value at these loci is lower when using the harmonized controls for GWAS instead of the internal controls. These loci provide examples of regions where there is a boost in power to detect true signals of association when using the harmonized controls.

**Figure 3.**
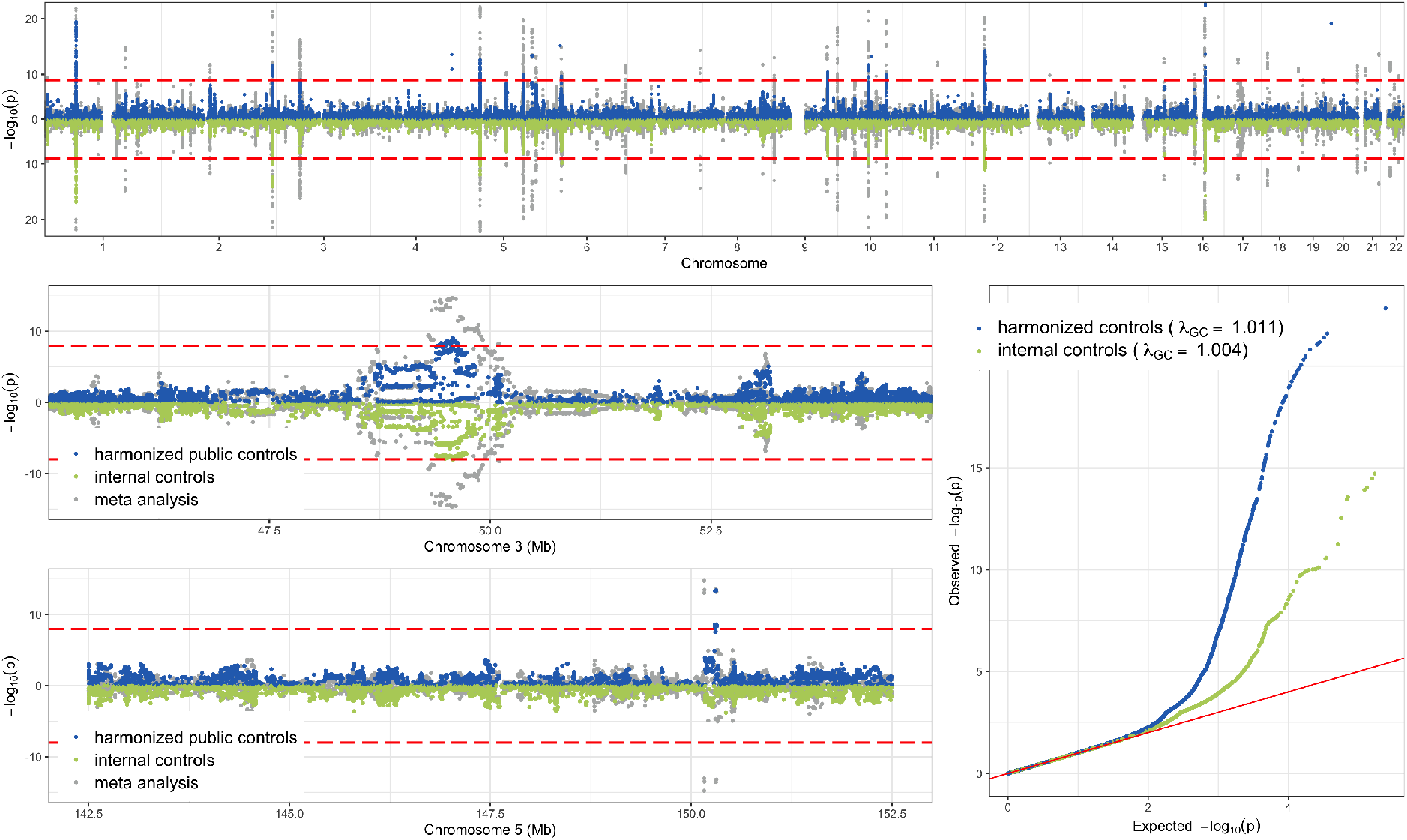
Comparison of GWAS for the Crohn’s disease study using harmonized public controls versus internal controls. Top: whole genome Manhattan plot. Bottom left: zoom-in to regions in individual chromosomes. Bottom right: QQ-plot of p-values (with respect to theoretical null).

Next, with the meta analysis as our best estimate of the truth, we directly compare the p-values obtained in GWAS (i) and (ii), examining separately SNPs that are significant and non-significant in meta-analysis (a p-value < 5 × 10^−8^ is considered significant). For variants that are not significant in the meta analysis (left panel of Figure 4), both GWAS determined these SNPs to be non-significantly associated with IBD as well, suggesting that the use of harmonized controls does not result in a higher false positive rate. On the right panel of Figure 4, looking at variants that are significant in meta-analysis, the majority of these sites are more significant in the GWAS of harmonized controls than in the GWAS of the internal controls. In particular, there are a number of variants that are deemed significant using the harmonized controls but missed using the internal controls. These results confirm that the use of harmonized controls yields no p-value inflation at null SNPs and boosts power at signal SNPs.

**Figure 4.**
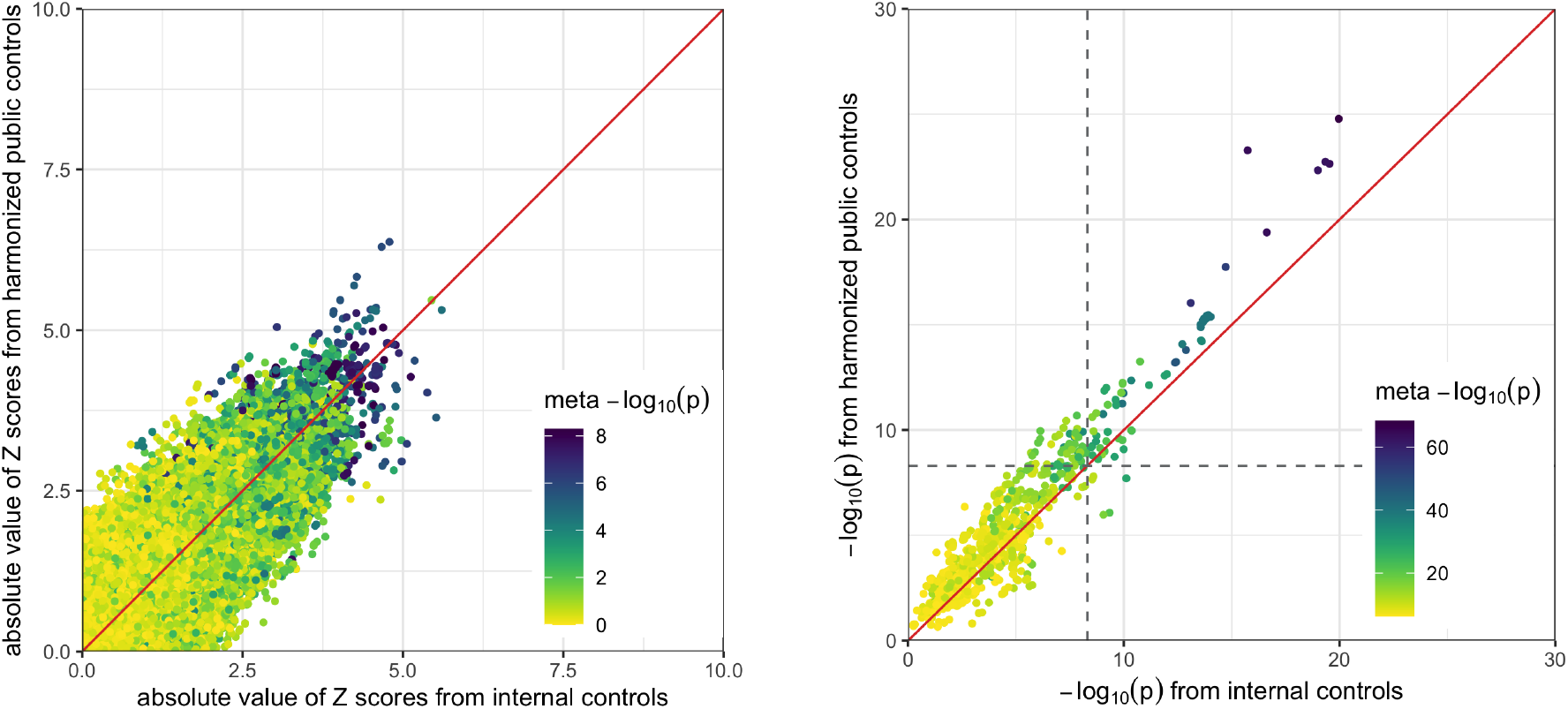
Comparison of p-values from using harmonized controls with those from using internal controls. Left: SNPs with p-values ≥ 5 × 10^−8^ in meta-analysis (for a better visualization, we plot the absolute Z-scores). Right: SNPs with p-values < 5 × 10^−8^ in meta-analysis (the dashed lines correspond to 5 × 10^−8^).

We next show examples of the ability of the harmonization pipeline to detect and correct for spurious associations that are driven by poor quality imputation, that otherwise would not be filtered in a standard GWAS pipeline. Figure 5 shows the Manhattan plots of the association results of analyses (i) and (iii) for two 20 Mb regions of Chromosome 1 (top row) and Chromosome 2 (bottom row). The left two plots show the results from GWAS (i) both before and after applying the *Re-imputation* Module 4, against the background of those from the meta analysis (GWAS (iii)) shown in grey. The red points correspond to genotyped variants with EmpRsq < 0.1. A large fraction of these red variants indeed generate spurious signals in the surrounding region in the first round of imputation: the p-values based on aggregated controls (before re-imputation) is small, but the meta-analysis suggests that they are not true signals. Comparing the lower and upper halves of the Manhattan plots we can see that removal and re-imputation of these red SNPs removes the spurious peaks around those points (these peaks were the consequence of poor imputation driven by the red variants). In the right two plots, we compare p-values calculated from the harmonized controls (GWAS (i)) with p-values from internal controls (GWAS (ii)) and observe that they are highly concordant. These results suggest that the *Re-imputation* module is effective in removing spurious signals that would otherwise appear when using the harmonized controls.

**Figure 5.**
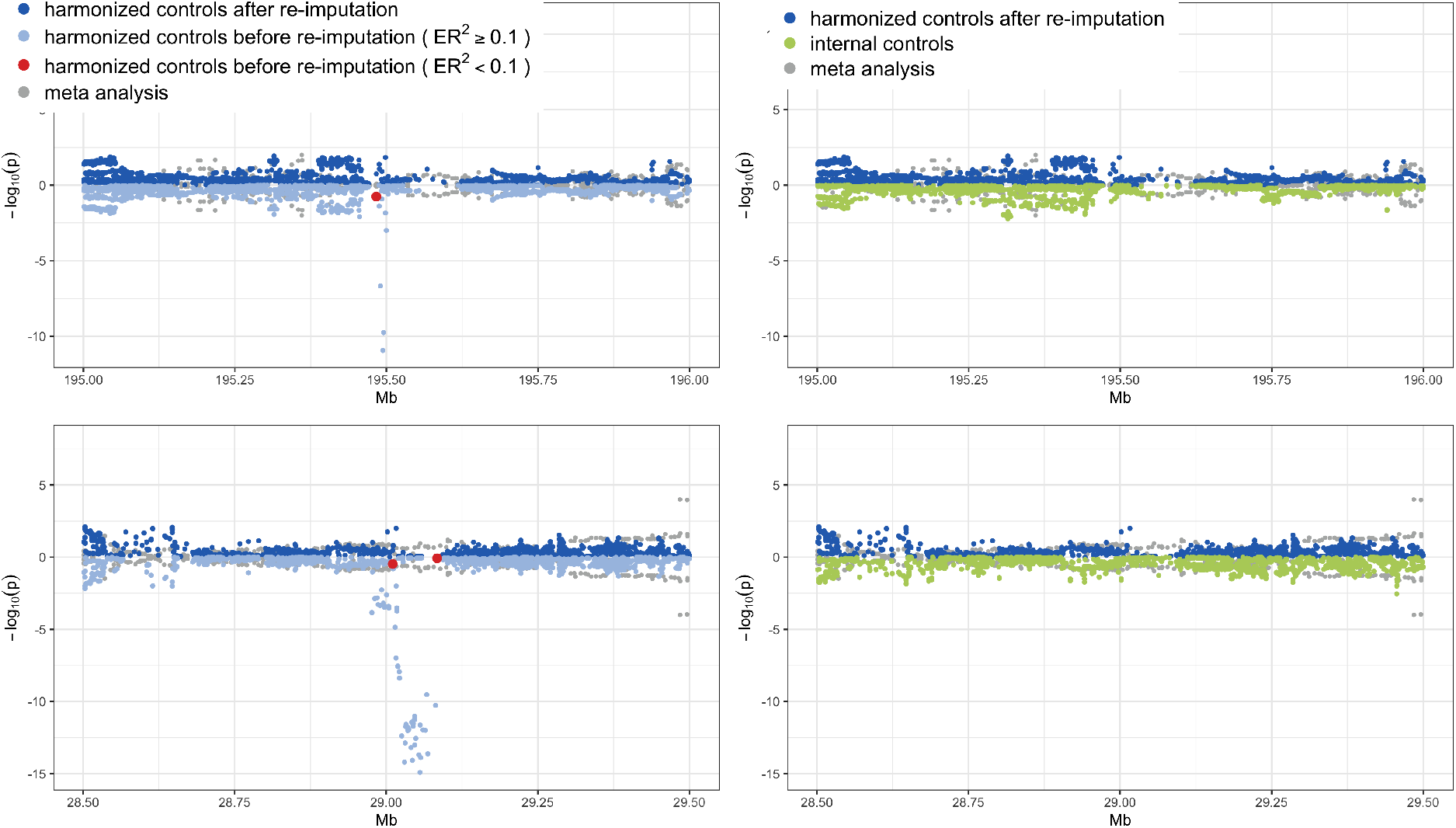
Examples of spurious signals being removed in the harmonization pipeline (top: Chromosome 1; bottom: Chromosome 2). The red dots are typed variants that drive poor quality imputation, flagged by their Em-pRsq values, which co-localize with batch effects. These spurious signals are corrected by re-imputation.

**Figure 6.**
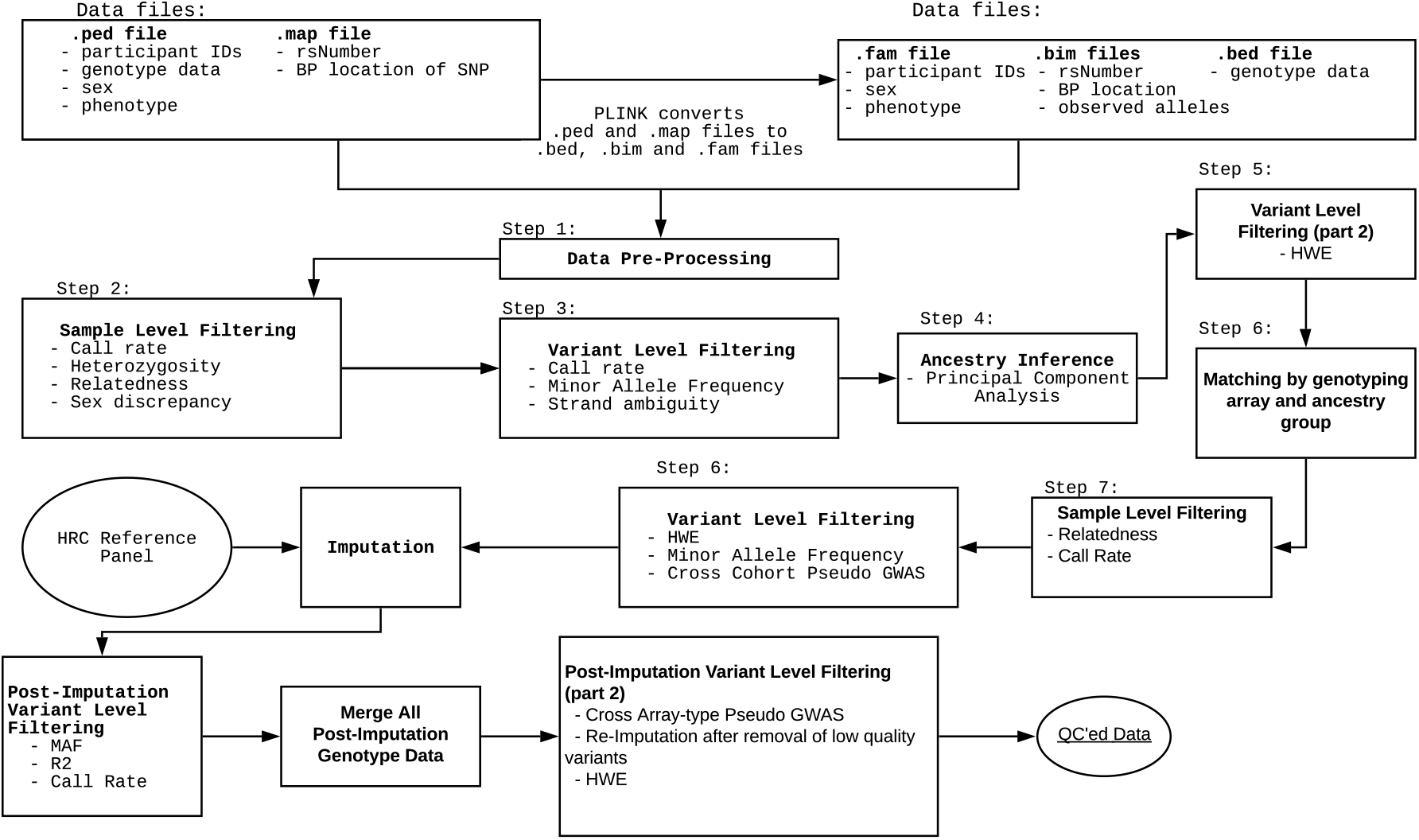
The complete QC Pipeline

## 4. Discussion

We have shown that it is possible to aggregate disparate genotyping data sets, even those assayed using different genotyping arrays, through a harmonization pipeline that involves iterative QC and imputation steps to control for batch effects and technical artifacts. This approach is valuable in constructing a large harmonized data set of external controls for use in GWAS, and we have shown it can deliver more powerful association tests while being robust to spurious signals driven by batch effects or insufficient ancestry matching.

The strength of our pipeline comes from the thorough and agglomerative approach to QC, that first operates within an array type for a single ancestry, and then across different arrays. The identification of problematic SNPs (through the EmpRsq metric) that are driving poor quality imputation, and the re-imputation after removing these sites is a key insight that allows the aggregation across different array types. Since multiple genotyping arrays have been used in human genetic studies, and as new arrays are developed over time, this step is essential to bring together different datasets from various sources.

While our approach permits existing external controls to be integrated in to GWAS, the extent to which these resources can be useful depends upon whether the ancestry spanned by the control set sufficiently covers that of the cases to which it is being compared. That is, for a case collection from a population that is underrepresented in publicly available controls, there will be a paucity of control samples of matched ancestry. If the harmonized control set does not sufficiently capture the ancestry space spanned by the cases, then the projection of cases on to the control-generated PC space will be biased^25^, giving an inaccurate depiction of their ancestry relative to the axes spanned by the control set. This mismatching of controls to cases can lead to spurious associations and subsequent false findings, and highlights the unmet need for the inclusion of more ancestrally diverse samples in public genetic resources.

We have not examined the application of this method to admixed samples, which pose a challenge as their ancestry is heterogeneous across the genome. This means that clustering individuals by PCA will group those that share similar proportions of ancestry genome-wide, and will not necessarily match samples by their ancestral origins at any specific genomic site. We propose the extension of this method to admixed populations as a future research direction.

Although we have demonstrated that it is possible to leverage multiple different sources of genotyping data for their use in a unified GWAS, the extensive quality control pipeline that we developed speaks to the many challenges and potential pitfalls involved in this process. We advocate for broad consent and data use agreements that enable public sharing of individual-level data to enable direct GWAS for health-related research. As public data sets grow increasingly larger, through biobanking efforts for examples, there will be less need to harmonize between different sources of control samples. Until then, the careful aggregation of multiple smaller resources can be valuable in enabling well powered GWAS at no additional cost.

## 5. Author contributions

D.C., A.B., C.C., K.T., D.P. designed the data harmonization pipeline. D.C. implemented the pipeline, performed all analyses and produced the figures. K.T. conducted the imputation. A.B., K.R. and B.N. supervised the project. Z.T.K. and C.C. wrote the manuscript and several authors provided valuable edits.

## Appendix A. 16 Genotype data collections in dbGap used to construct the harmonized control dataset

In Section 3.1, we create a resource of external controls using genotype data taken from 27,517 individuals of self-reported European-descent in 16 studies in the dbGaP repository. Information regarding these cohorts can be found in Table 2.

## Appendix B. The complete QC Pipeline for data harmonization

We give a step-by-step description of the QC pipeline. First, we conducted pre-processing:

(1) All cohorts were lifted over from hg18 to hg19 using the liftOver tool. Marker names across all cohorts were made consistent with the HRC reference panel based on chromosome, position, and alleles. Indels and missing alleles, duplicated markers, and non-matching SNPs were excluded.

Next, we conducted cohort-level pre-imputation QC:

(2) *Sample QC:* samples with missing rate greater than 2% or with mismatching sex information between genotypes and provided phenotype information were excluded. Samples with abnormal inbreeding coefficients (3 standard deviations away from the mean) were removed. Related samples (pi-hat > 0.625) were identified and the sample in each related pair with higher missingness was removed from the dataset.
(3) *Variant QC:* variant-level missingness was calculated and variants with missing rate > 1% were excluded. Additionally, all A/T and C/G SNPs were excluded at this point to avoid downstream issues related to strandness.
(4) *Ancestry Inference:* we built a random forest classifier (see below for details) with the first six PCs from 1000 genomes to determine broad ancestry grouping. We then further separate population isolates from major European within the European ancestry cluster by building a second random forest classifier using the first four PCs calculated from a merged set of the 1000 genomes Europeans and controls from an Ashkenazi Jewish cohort. We then projected each sample onto the 1000 genomes PC space, and assigned broad ancestry-group labels using the first random forest classifier. Each sample was assigned to one continental ancestry group. A similar procedure was applied to samples assigned a European ancestry label to determine its finer ancestral origin. Within each ancestral group, variants with minor allele frequencies lower than 0.01 or with p-values for Hardy-Weinberg Equilibrium test less than 10^−4^ were excluded. Random forest classifier details: We used the following Random Forest algorithm to perform ancestry assignment. We first assigned individuals into one of five super-populations: East Asian, South Asian, African, American, and European. To do so, we first calculated the PCs of the 1000 Genomes dataset. We then apply *RandomForestClassifier* from *scipy* package and train a random forest classifier with 100 bootstrap draws using the first six PCs as input. All parameters of the random forest training function were set at their defaults. We then used the output classifier to predict the population label of each individual. For individuals that are assigned as European, we further assigned them into major European Ancestry, Finnish Ancestry and Ashkenazi Jewish Ancestry. To do this, we merged 1000 Genomes European genotype data with genotype data from an Ashkenazi Jewish cohort as there are no Ashkenazi Jewish individuals among the 1000 genomes samples. We then evaluate PCs in this merged dataset. Non-Finnish Europeans are assigned the *Major European* population group label. The first four PCs are used to train another the random forest model. Again, all parameters of the random forest training function in *RandomForestClassifier* are set at their defaults. The resultant classifier was used to assign finer structure in the European subset. ^26^.

Next, we conducted array-level pre-imputation QC:

(5) *Matching:* cohorts from the same genotyping array and ancestry-group were merged together for downstream analysis.
(6) *Sample QC:* samples with missing rate greater than 2% and the member of related pairs (pi-hat > 0.625) of samples with higher missingness was removed.
(7) *Variant QC:* within each array group, variants with minor allele frequencies lower than 0.01 or with p-values for a Hardy-Weinberg Equilibrium test less than 10^−4^ were excluded. To further identify variants susceptible to batch effects, a pseudo-GWAS comparison labelling array samples as cases and 1000 Genomes samples as controls was performed. Variants with p-value less than 10^−4^ were excluded. Further pseudo-GWASes defeined by labelling samples from one cohort as cases and samples from all other cohorts as controls were carried out, and variants with p-values less than 10^−4^ were also excluded. For pseudo-GWAS comparison, the first 20 PCs were included as covariates to control for population stratification.

The next step is imputation:

(8) We imputed each stratum separately using 1000 genomes as a reference panel via the Michigan Imputation server.

Then, we have a few steps of post-imputation QC:

(9) *QC each array stratum:* After imputation, 46.8M variants had been imputed in each array stratum. Variants with minor allele frequencies < 0.01, out of Hardy-Weinberg Equilibrium (p < 10^−4^) or with an imputation info score < 0.8 were removed from the data.
(10) *Inner-join:* a master imputed genotype dataset was generated through an inner join across all stratum.
(11) *Cross-array comparison:* for each genotyping array, samples that were genotyped on that array were coded as cases and samples that were genotyped on any other array were coded as controls to make a pseudo-case control comparison. To run association testing, we added first 20 PCs as covariates and drop variants found to be associated with any given array with p-value < 10^−5^.
(12) *Re-imputation:* We removed variants with empRsq < 0.6 from the typed data, and re-imputed them back so as to increase the number of SNPs as well as to improve the quality. The re-imputation procedure repeats steps (8)-(11).

## Appendix C. The QC Pipeline on cohron’s disease data

We conducted GWAS on five Crohn’s disease genetics data sets from the IBD Genetics Consortium, to evaluate the quality of the collection of harmonized controls. We applied a QC pipeline to these five data sets. This QC pipeline is a subset of the data harmonization pipeline in Appendix B, where we skip the steps for cross-array comparison and re-imputation.

We now describe the QC pipeline on Crohn’s disease data.

- We first conducted data pre-processing to align cases to hg19 Genome Build. This is the same as step (1) of the data harmonization pipeline in Appendix B.
- Next, we conducted cohort-level pre-imputation QC:
  – *Sample QC:* We filtered out samples with high missing rate, mismatching sex, abnormal inbreeding coefficients and related individuals. This is the same as step (2) of the data harmonization pipeline.
  – *Variant QC:* We filtered out high missing rate and strand ambiguous SNPs. This is the same as step (3) of the data harmonization pipeline.
  – *Ancestry Inference:* In step (4) of the data harmonization pipeline, we built a Random Forest Classifier for classifying any sample into one of the five super populations: East Asian, South Asian, African, American, and European. We applied this classifier to assign each sample into a super population. In step (4) of the data harmonization pipeline, we also built a Random Forest Classifier to classify any European sample into one of the three groups, European Mainland, Finnish, and Ashkenazi Jewish. We applied this classifier to further divide those samples assigned to the European super population.
- We then imputed case cohort against 1000 Genomes panel. This is the same as step (8) of the data harmonization pipeline.
- Lastly, we performed post-imputation QC: We remove variants out of Hardy-Weinberg Equilibrium with low minor alleles frequency, or with low imputation INFO score. This is the same as step (9) of the data harmonization pipeline.

After the above QC pipeline is completed, if the GWAS results still have many suspicious signals, the user can add a further re-imputation step, the same as step (12) in the data harmonization pipeline. In the case of the Crohn’s disease data, we do not require reimputation.

## Notes

https://github.com/mikkoch/unicorn-qc

### Competing Interest Statement

The authors have declared no competing interest.

